# CLASP2 facilitates dynamic actin filament organization along the microtubule lattice

**DOI:** 10.1101/2022.09.22.509100

**Authors:** NC Rodgers, EJ Lawrence, AV Sawant, N Efimova, G Gonzalez-Vasquez, TT Hickman, I Kaverina, M Zanic

## Abstract

Coordination between the microtubule and actin networks is essential for cell motility, neuronal growth cone guidance, and wound healing. Members of the CLASP (Cytoplasmic Linker-Associated Protein) family of proteins have been implicated in the cytoskeletal crosstalk between microtubules and actin networks, however, the molecular mechanisms underlying CLASPs role in cytoskeletal coordination are unclear. Here, we investigate CLASP2α’s crosslinking function with microtubules and F-actin. Our results demonstrate that CLASP2α crosslinks F-actin to the microtubule lattice in vitro. We find that the crosslinking ability is retained by L-TOG2-S, a minimal construct containing the TOG2 domain and serine-arginine rich region of CLASP2α. Furthermore, CLASP2α promotes the accumulation of multiple actin filaments along the microtubule, supporting up to 11 F-actin landing events on a single microtubule lattice region. CLASP2α also facilitates dynamic organization of polymerizing actin filaments templated by the microtubule network, with F-actin forming bridges between individual microtubules. Finally, we find that depletion of CLASPs in vascular smooth muscle cells results in disorganized actin fibers and reduced co-alignment of actin fibers with microtubules, suggesting that CLASP and microtubules contribute to higher-order actin structures. Taken together, our results indicate that CLASP2α can directly crosslink F-actin to microtubules, and that this microtubule-CLASP-actin interaction may influence overall cytoskeletal organization in cells.

## INTRODUCTION

The cytoskeleton is an essential cellular component that drives a multitude of processes such as cell division and motility, and defines cell shape. Individual components of the cytoskeleton must coordinate to perform their cellular functions (Dogterom & Koenderink, 2019; Pimm & Henty-Ridilla, 2021; Rodriguez et al., 2003). For example, microtubules and actin interact with each other to facilitate cell motility (Ballestrem et al., 2000; Salmon et al., 2002; Waterman-Storer & Salmon, 1999, 1997; X. Wu et al., 2008) and growth cone guidance (Dent & Kalil, 2001; Rochlin et al., 1999; Schaefer et al., 2002; Slater et al., 2019). Many cellular factors are involved in this coordination (Rodriguez et al., 2003), and the proteins that physically couple the microtubule and actin networks are of particular interest (Dogterom & Koenderink, 2019; Pimm & Henty-Ridilla, 2021). However, the specific interactions underlying cytoskeletal coupling remain to be fully understood.

Cytoplasmic linker associated proteins (CLASPs) are a well-studied family of microtubule-associated proteins (MAPs), that have been implicated in interacting with both microtubules and F-actin (Dogterom & Koenderink, 2019; Engel et al., 2014; Tsvetkov et al., 2007). CLASPs are known as microtubule stabilizers with essential roles in cell division, cell migration and neuronal development (Lawrence et al., 2020). There are two paralogs of CLASP, CLASP1 and CLASP2, that are alternatively spliced resulting in multiple isoforms, which are differentially expressed and may have some isoform-specific functions (Akhmanova et al., 2001; Lawrence et al., 2020). In vitro studies with purified proteins established that CLASPs promote sustained microtubule growth by suppressing microtubule catastrophe, the transition from growth to shrinkage, while promoting microtubule rescue, the transition from shrinkage to growth (Aher et al., 2018; Akhmanova et al., 2001; Al-Bassam et al., 2010; Lawrence et al., 2018; Leano et al., 2013; Maki et al., 2015; Patel et al., 2012). In cells, CLASPs are targeted to growing microtubule plus ends via a direct interaction with microtubule end-binding EB proteins and can specifically regulate microtubule dynamics at the actin-rich cell cortex (Mimori-Kiyosue et al., 2005). Furthermore, CLASPs are involved in the process of microtubule plus-end capture at the cell cortex through interactions with LL5β, a component of the cortical microtubule stabilization complex (Hotta et al., 2010; Lansbergen et al., 2006; Stehbens et al., 2014). CLASPs’ stabilization and anchoring of microtubules is also important for the regulation and dynamics of podosomes, polymerizing actin-based structures, in vascular smooth muscle cells (Efimova et al., 2014; Zhu et al., 2016). All these studies suggest CLASPs’ roles in the interplay between the microtubule and actin networks. However, it remains unclear if CLASPs directly interact with the actin network in these contexts.

A previous study directly investigating CLASP-actin interaction reported that the two CLASP paralogs, CLASP1 and CLASP2, colocalize with actin stress fibers in primary fibroblasts and spinal cord neurons (Tsvetkov et al., 2007). The authors found that both CLASP paralogs coimmunoprecipitated with actin from fibroblast cells and suggested that the tumor overexpressing gene (TOG)-1 domain and serine-arginine rich region of CLASP2α facilitate the F-actin interaction. Another study, using co-sedimentation experiments with purified proteins reported a direct interaction between F-actin and CLASP2α, as well as co-sedimentation of G-actin with microtubules in the presence of CLASP2α (Engel et al., 2014). To our knowledge, these two reports provide the only evidence of CLASP-actin interaction, leaving open the question of whether CLASP alone is sufficient for crosslinking of microtubules and actin filaments, which may play a role in the organization of stress fibers in cells.

## RESULTS AND DISCUSSION

### Human CLASP2α directly crosslinks actin filaments to microtubules

To investigate CLASP2α’s ability to directly crosslink microtubules and actin, we expressed and purified full-length CLASP2α using an Sf9-insect-cell based system (Figure 1A and Supplemental Figure S1A). Performing co-sedimentation experiments with purified CLASP2α and polymerized purified F-actin, we found a statistically significant increase in the amount of CLASP2α in the pellet (Supplemental Figure S1A). Thus, consistent with previous reports (Engel et al., 2014), we find that CLASP2α directly interacts with F-actin.

**Figure 1.**
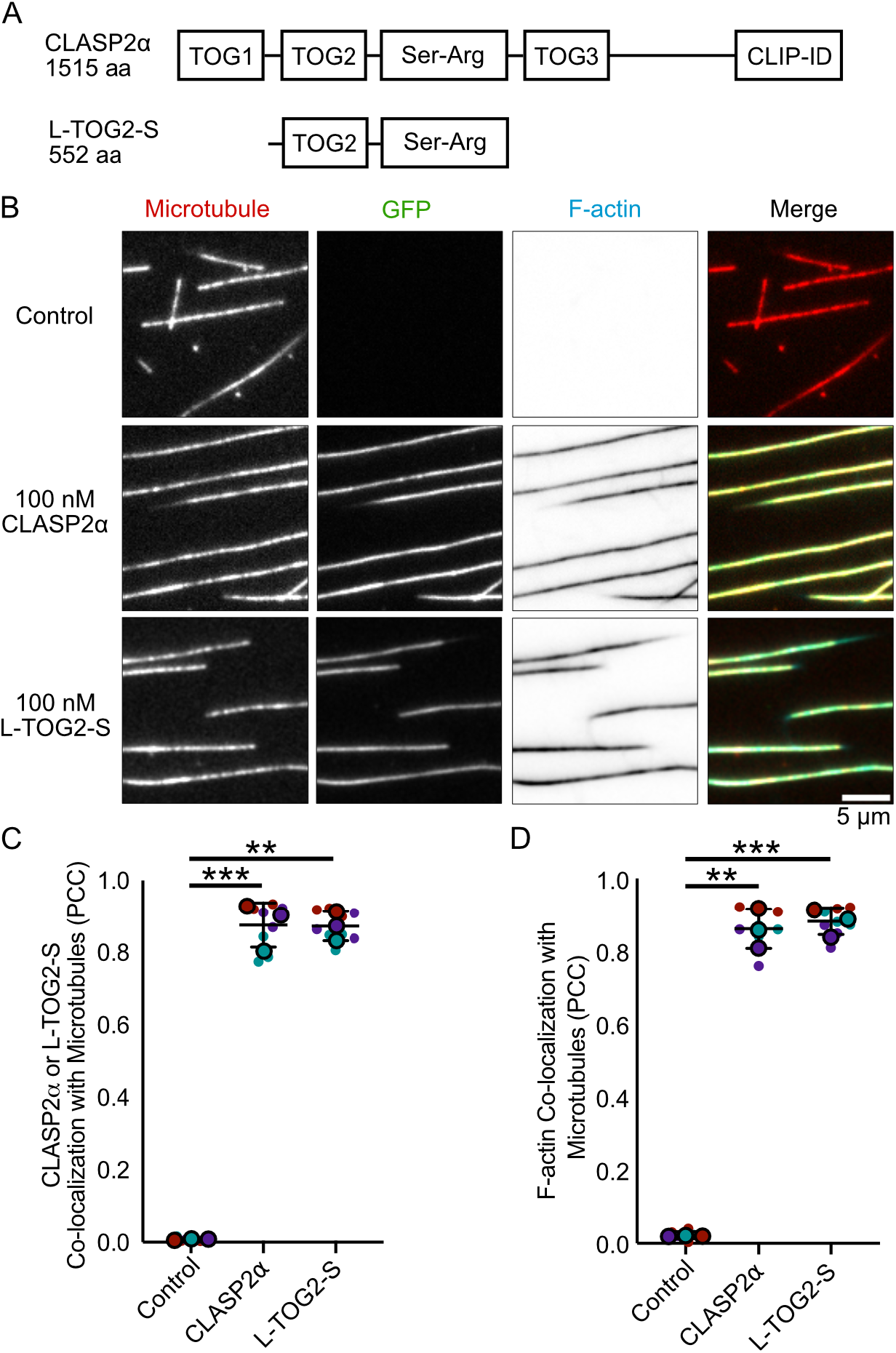
Human CLASP2α directly crosslinks actin filaments to microtubules. A) Schematic of the domain structure of human CLASP2α and minimal construct, L-TOG2-S. B) Representative images of Taxol-stabilized microtubules incubated for 10 minutes with storage buffer, 100 nM CLASP2α-GFP, or 100 nM GFP-L-TOG2-S and 6.5 μM TRITC-Phalloidin F-actin. C) Quantification of co-localization between either CLASP2α or L-TOG2-S with microtubules using the Pearson correlation coefficient (PCC). D) Quantification of co-localization between actin filaments and microtubules, using the PCC. Small data points are individual PCC measurements and large data points are the mean PCC for three independent experimental days, represented by color. Error bars are the standard deviation. Kruskal-Wallis test multiple comparisons, ** p < 0.01 and *** p < 0.001. Comparisons between CLASP2α and L-TOG2-S are ns, p > 0.05.

To determine if CLASP2α can simultaneously bind to microtubules and F-actin, we employed an *in vitro* reconstitution assay using 3-color imaging by total internal reflection fluorescence (TIRF) microscopy. First, we bound Alexa-647-labeled, Taxol-stabilized microtubules to the surface of a coverslip. Next, we added 6.5 μM TRITC-phalloidin-stabilized F-actin with or without 100 nM purified CLASP2α-GFP (Supplemental Figure S1B) to the flow-cell. As expected, we observed specific CLASP2α-GFP localization along the microtubule lattice (Figure 1B). We quantified the correlation between the CLASP2α-GFP and microtubule signals after 10 minutes of incubation using the Pearson Correlation Coefficient (PCC) and measured a high degree of correlation (PCC = 0.88 ± 0.06, SD, N = 9 fields of view (FOV) over 3 independent experimental days) (Figure 1C). The investigation of TRITC-phalloidin-F-actin showed that F-actin robustly co-localized with microtubules in the presence of CLASP2α-GFP (PCC = 0.86 ± 0.05, SD, N = 9 FOVs over 3 independent experimental days) (Figure 1B,D). No significant co-localization between microtubules and F-actin was observed in the absence of CLASP2α (PCC = 0.022 ± 0.001, SD, N = 9 FOVs over 3 independent experimental days) (Figure 1B,D). Furthermore, we confirmed that the co-localization of F-actin with microtubules does not depend on phalloidin-actin stabilization, as F-actin also localized to CLASP2-coated microtubules without phalloidin (Supplemental Figure S2A). Taken together, we conclude that CLASP2α directly crosslinks F-actin to microtubules.

Previous reports using co-immunoprecipitation from fibroblast cells implicated a Serine-Arginine rich region of CLASP2α in its interaction with actin (Tsvetkov et al., 2007). Recently, a minimal CLASP2 construct, L-TOG2-S, containing a single TOG2 domain and the Serine-Arginine rich region (Figure 1A), was reported to recapitulate CLASP’s effect on microtubule dynamics (Aher et al., 2018). We wondered whether this minimal CLASP2 construct is also sufficient to crosslink microtubules and F-actin. We used the same TIRF-based approach to investigate TRITC-phalloidin-F-actin localization on microtubules in the presence of purified 100 nM GFP-L-TOG2-S (Supplemental Figure S1B). As expected, GFP-L-TOG2-S showed strong localization to microtubules (PCC = 0.87 ± 0.04, SD, N = 9 FOVs over 3 independent experimental days) (Figure 1B-C). Furthermore, we found that F-actin also robustly co-localized with microtubules in the presence of GFP-L-TOG2-S (PCC = 0.88 ± 0.04, SD, N = 9 FOVs over 3 independent experimental days) (Figure 1B,D). This result demonstrates that the minimal CLASP2α construct containing a single TOG2 domain, and the Serine-Arginine rich region is sufficient to directly crosslink F-actin to microtubules.

### CLASP2α mediates sequential binding of actin filaments along the microtubule lattice

To further investigate how actin filaments interact with microtubules in the presence of CLASP2α, we imaged TRITC-phalloidin-F-actin on microtubules over time for up to 40 minutes. We observed the initial landing of actin filaments within two minutes (Figure 2A, Video 1). Notably, the F-actin signal continued to increase over time, reaching saturation within the duration of the experiment (Figure 2B). No F-actin binding to microtubules was observed in the absence of CLASP2α for the entire duration of the experiment (Figure 2A, Video 1).

**Figure 2.**
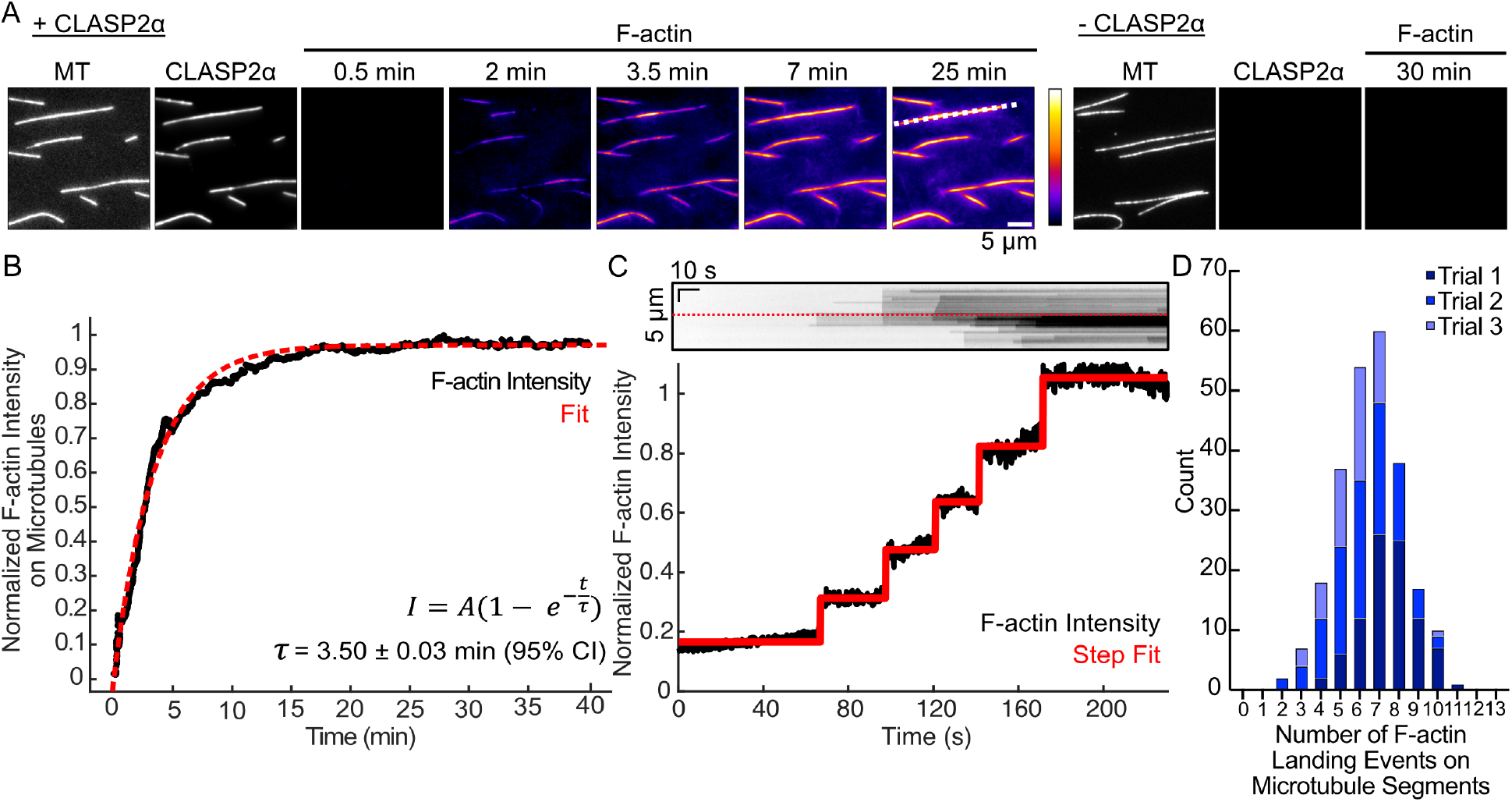
CLASP2 mediates sequential binding of actin filaments along the microtubule lattice. A) Example time-lapse TIRFM images of Taxol-stabilized microtubules, in the presence of 100 nM CLASP2α-GFP or CLASP2α storage buffer, and addition of 6.5 μM TRITC-phalloidin-stabilized F-actin. White dotted line corresponds to the kymograph in Panel C. B) Example normalized actin filament intensity in microtubule areas over time for the images in panel A. Red dotted line represents the fit to the intensity over time equation. C) Top: kymograph of the Factin channel for an example microtubule (denoted by a dotted orange line in Panel A) for the first 5 minutes. Red dotted line corresponds to the intensity line scan. Bottom: 3-pixel intensity line scan for the F-actin channel in the kymograph. Red line represents the vbFRET stepping algorithm fit to count steps of F-actin landing events (see Methods). D) Stacked histogram of the total number of F-actin landing events on microtubule segments. Experiments done in triplicate (N = 91 (Trial 1), 99 (Trial 2), and 54 (Trial 3) microtubule regions analyzed).

A closer inspection of the increasing TRITC-phalloidin-F-actin intensity on individual microtubules revealed step-like increases in fluorescence intensity on a region of microtubule lattice (Figure 2C), suggesting sequential binding of actin filaments. Using a stepping analysis (Bronson et al., 2009, see Methods), we measured the accumulation of F-actin within a 3-pixel-wide (480 nm) microtubule segment over a period of 40 minutes. The mean number of landing events was 6.3 ± 0.6 (SD, N = 91, 99, and 54 microtubule lattice segments analyzed on three experimental days), with up to 11 sequential F-actin landing events occurring on a single microtubule segment (Figure 2D).

We wondered if the sequential accumulation of F-actin on microtubules could be supported by potential F-actin bundling activity of CLASP2α. However, using a low-speed co-sedimentation approach, we did not find any evidence that CLASP2α can bundle F-actin (Supplemental Figure S2B-D). In these experiments, bundled F-actin preferentially sediments into the pellet, while single F-actin filaments remain in the supernatant, due to low centrifugation speed. As a positive control, we used α-actinin, a well-known actin bundler, and observed a significant increase in the amount of F-actin in the pellet (Supplemental Figure S2C,D). In contrast, no significant increase in the F-actin pellet fraction was observed in the presence of CLASP2α (Supplemental Figure S2B,D). This result suggests that the CLASP2α-mediated F-actin accumulation on microtubules does not occur through bundling of actin filaments by CLASP2α.

### CLASP2α facilitates dynamic actin filament organization templated by the microtubule arrangement

We next wondered if CLASP2α could facilitate dynamic actin polymerization along the microtubule lattice. To probe this, we introduced 250 nM soluble, Alexa-647-labeled, globular actin (G-actin) into a flow-cell containing Taxol-stabilized, coverslip-bound microtubules (Figure 3). In the presence of CLASP2α, we observed the binding and growth of dynamic actin filaments along the microtubule lattice, which continued to grow off the ends of the microtubule polymer. (Figure 3A,B). Although individual microtubules were sparsely distributed on the coverslip surface, we often observed growing F-actin forming connections between microtubule polymers (Figure 3A, Video 2). We measured the length of the individual F-actin connections between microtubules to be 7 ± 3 μm (SD, N = 26 F-actin bridges over 3 independent experimental days, Figure 3C). Thus, under these conditions, CLASP2α can promote linking of microtubules by actin filaments. Without CLASP2α, microtubules were not covered by F-actin and no connections were observed (Figure 3A). These results demonstrate that CLASP2α can support dynamic F-actin organization templated by microtubules, and that F-actin can bridge microtubules forming an interconnected network.

**Figure 3.**
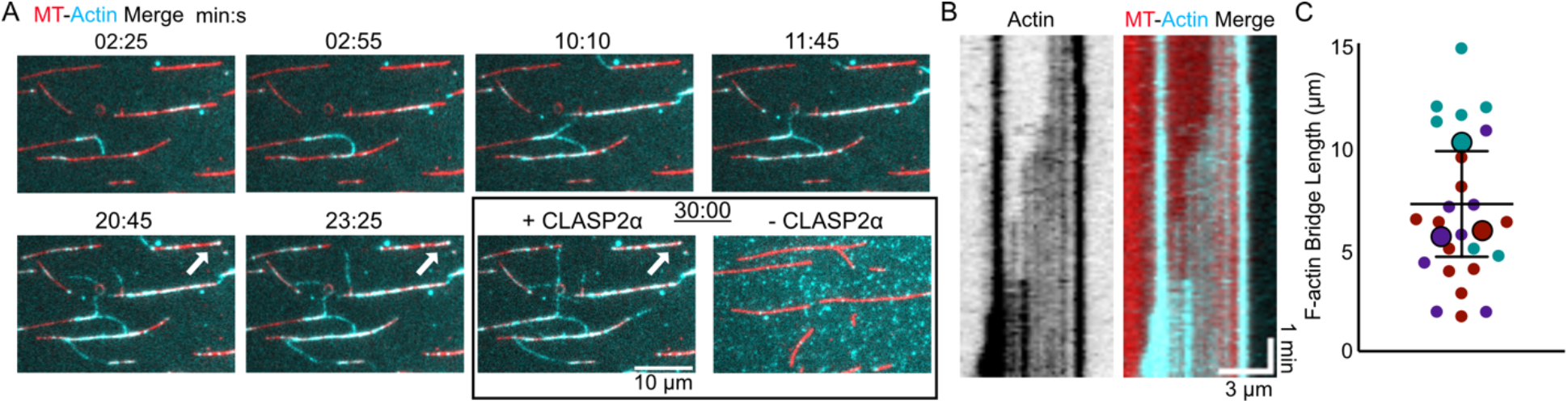
CLASP2 facilitates dynamic actin filament organization templated by the microtubule network. A) Example time-lapse images demonstrating dynamic F-actin (cyan) connecting multiple microtubules (red). White arrow highlights an example of growing F-actin shown by a kymograph in Panel B. Last image represents the buffer control (-CLASP2α) after 30 minutes. B) Example kymograph demonstrating growing F-actin along a microtubule lattice (left: actin channel alone, right: merged image). C) Quantification of the individual lengths of F-actin connections between microtubules. Smaller data points are the individual measurements and larger data points are the mean lengths for each repeat (N = 12, 7, and 7). The error bar is the standard deviation of the mean lengths.

### CLASP depletion results in disorganized actin fibers in cultured vascular smooth muscle cells

To probe CLASPs’ role in actin organization in cells, we depleted CLASPs in a rat vascular smooth muscle cell line A7r5, chosen due to its notable actin bundle (stress fiber) structures (Figure 4A). For prominent phenotypes, both CLASP paralogs (CLASP1 and CLASP2) were depleted by siRNA oligo combinations, carefully validated in previous work (Efimova et al., 2014; Zhu et al., 2016), achieving reliable reduction of CLASP protein levels (Supplemental Figure S3). Our results showed that actin fiber organization was severely disturbed in CLASP-depleted cells (Figure 4B,C). Actin organization was efficiently rescued by ectopic overexpression of non-silenceable mutant of CLASP2 (Fig. 4D) in depleted cells, indicating that CLASP2 paralog is likely sufficient for proper actin organization in this context. Interestingly, co-alignment of actin fibers with microtubules in CLASP-depleted cells (Figure 4F,G) was reduced as compared to controls (Figure 4E), suggesting a diminished coordination of these filaments, consistent with our *in vitro* results. Overall, our findings suggest that CLASPs contribute to co-organization of higher-order actin structures with microtubules in cells.

**Figure 4.**
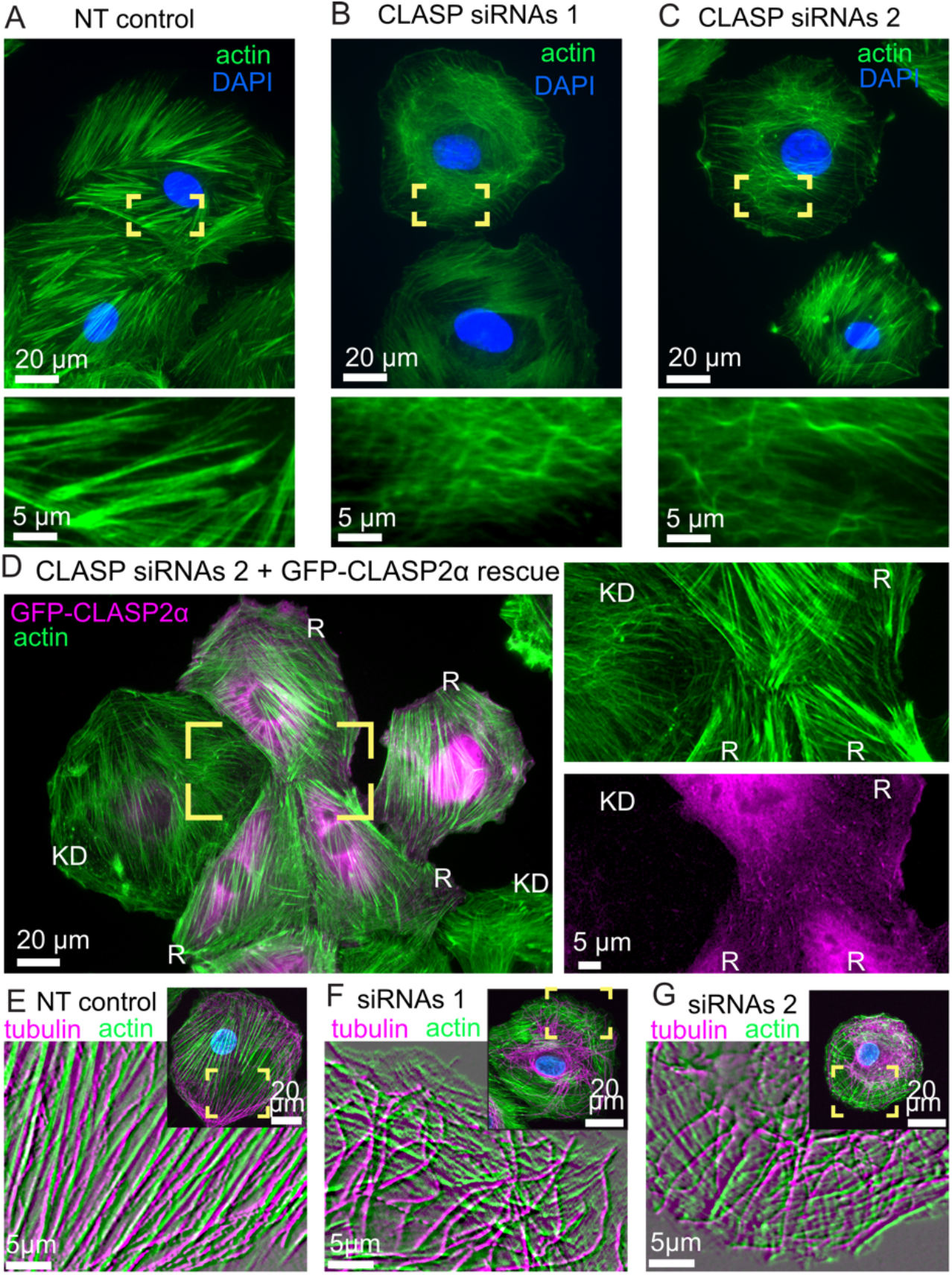
CLASPs are essential for correct stress fiber organization in vascular smooth muscle cells. A) Prominent actin stress fibers in cells treated with a control (non-targeting, NT) siRNA. B-C) Disorganized actin mesh in cells treated with two alternative combinations of siRNA oligos. Phalloidin-stained actin, green. DAPI, blue. Yellow boxes are enlarged below to highlight details. The phenotypes have been vetted by a double-blinded evaluation (see Methods). D) Cells treated with siRNA combination 2 and transfected with GFP-CLASP2 (pseudo-colored magenta) resistant to this siRNA. Separate channels from the yellow box are enlarged at the right. Actin organization in GFP-CLASP2-rescued cells (R) is similar to control (as in Panel A), while in a non-expressing cell (KD) it is similar to knockdown phenotype (as in Panel C). Phalloidin-stained actin, green. GFP-CLASP2, magenta. E-F) Actin and microtubules in cells treated with a control (non-targeting, NT) siRNA (E) or with siRNA combinations 1 (F) or 2 (G). Phalloidin-stained actin, green. Tubulin, magenta. DAPI, blue. Yellow boxes in overview images are enlarged in insets. Inset images are processed through the emboss filter for exclusive visualization of fibers and illustration of their alignment with microtubules, which is diminished in (F, G) as compared to (E). Scale bars, 20μm in overviews and 5μm in insets. All panels show representative images out of more than 45 cells per condition over 3 or more repeated experiments.

### Conclusions

In summary, our results demonstrate that CLASP2α directly crosslinks F-actin to microtubules. We find that a minimal CLASP construct, containing the TOG2 domain and the Serine-Arginine rich region of CLASP2, is sufficient to crosslink microtubules and F-actin (Figure 5). Interestingly, this L-TOG2-S construct was previously shown to recapitulate CLASP2’s effects on microtubule end dynamics (Aher et al., 2018). There, the authors reported that the TOG2 domain alone has a weak affinity for microtubules, and that CLASP’s unstructured, positively charged region is needed for robust direct microtubule binding. The same Serine-Arginine region contains a Ser-x-Ile-Pro (SxIP) motif, which encodes CLASP’s direct interaction with EB proteins, facilitating targeting of CLASP to growing microtubule ends (Maki et al., 2015; Patel et al., 2012). How interactions of the Serine-Arginine region with microtubules, actin and EBs are regulated in different cellular contexts presents an interesting question. A previous report using co-immunoprecipitation and co-localization experiments in fibroblasts suggested that the TOG1 domain of CLASP2α also interacts with F-actin (Tsvetkov et al., 2007). Our results show that the TOG1 domain is not necessary for the CLASP-mediated crosslinking of microtubules with F-actin; to what extent TOG1 domain may contribute to direct CLASP-actin interaction warrants further investigation. Interestingly, a recent report suggested that another TOG-domain protein, XMAP215, directly interacts with F-actin to promote microtubule-actin coalignment in the neuronal growth cone (Slater et al., 2019). There, the authors found that all five TOG domains of XMAP215 were required for XMAP215’s localization to F-actin. Future work will be needed to determine the affinities of individual TOG domains for F-actin and their effects on the microtubule-actin crosstalk.

**Figure 5.**
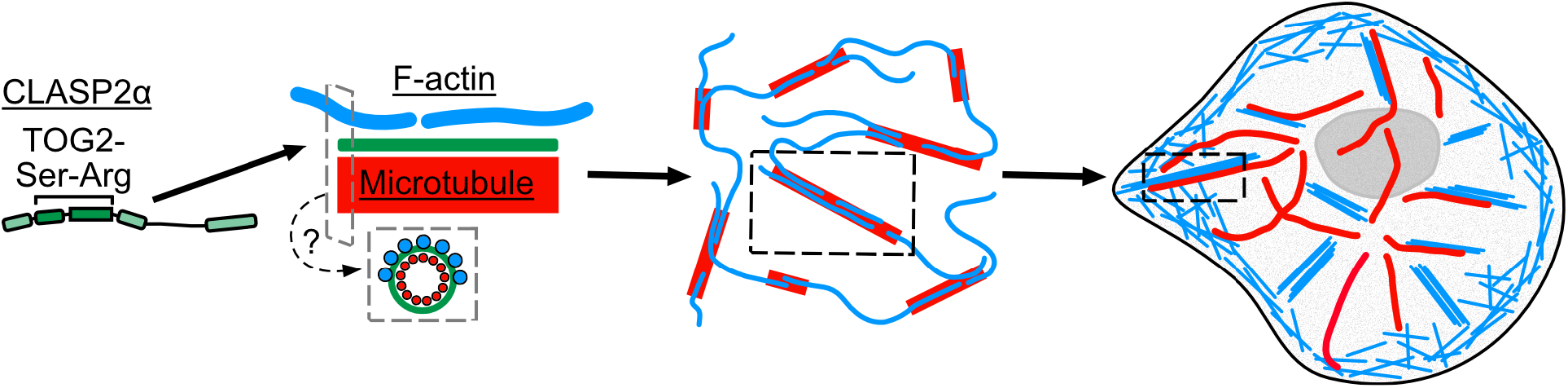
CLASP2 facilitates dynamic actin filament organization along the microtubule lattice. Microtubules are in red, actin is in blue, and CLASP2 is green throughout figure. Gray dashed boxes represent cross-sectional view of a potential model for F-actin organization along microtubules coated with CLASP2α, where F-actin are binding around the microtubule shaft. Black dashed boxes are highlighted zoom ins representing instances of co-organization of actin filaments and microtubules observed *in vitro* and in cells.

Our finding that microtubule-associated CLASP2α supports the binding of up to 11 actin filaments to a single microtubule region raises the question of F-actin organization on the microtubule lattice. Our results don’t show any evidence of actin forming bundles on the microtubule lattice. The number of F-actin landing events saturated within the duration of our experiment, and we never observed more than 11 sequential landing events, suggesting a limit to the number of F-actin that can be linked to the microtubule lattice. Furthermore, we found no evidence of CLASP2α being able to directly bundle F-actin. This is different from the effects of another well-studied MAP, tau, which directly bundles F-actin and facilitates actin elongation and bundling along growing microtubules (Elie et al., 2015). Given that Taxol-stabilized bovine microtubules typically contain 13 individual protofilaments (Amos & Klug, 1974; Arnal & Wade, 1995), and that the full microtubule surface may not be accessible to F-actin binding in our experiments (due to the tethering of microtubules to the coverslip), the number of landing events we observed is consistent with the model in which individual actin filaments bind around the microtubule perimeter along microtubule protofilaments (Figure 5). High resolution structural approaches, such as cryo-electron microscopy, will be needed to determine the exact organization of F-actin on CLASP2α-coated microtubules.

Our results demonstrate that CLASP2α-coated microtubules can template dynamic F-actin organization, specifically by F-actin linking multiple microtubules and forming an interconnected cytoskeletal network. Other crosslinking proteins have been shown to organize F-actin in a microtubule-centric fashion, with notable examples belonging to the spectraplakin protein family (Dogterom & Koenderink, 2019; Pimm & Henty-Ridilla, 2021; Rodriguez et al., 2003; Suozzi et al., 2012). Interestingly, a number of microtubule plus-tip interacting proteins (+TIP) have also been shown to interact with F-actin and influence actin dynamics. Cytoplasmic linker interacting protein 170 (CLIP-170), a prominent +TIP and a known binding partner of CLASP, was recently reported to directly bind to F-actin and microtubules, however the interaction of CLIP-170 with F-actin and microtubules was found to be mutually exclusive (Y.-F. O. Wu et al., 2021). Similarly, EB1 was reported to directly interact with F-actin, however it did not bind F-actin and microtubules simultaneously (Alberico et al., 2016). While some of the +TIPs might not be able to directly crosslink F-actin and microtubules, many interactions among +TIPs in combination with other actin binding proteins result in the collective regulation of microtubule-actin crosstalk. For example, the combination of CLIP-170 and EB1 with actin-regulators formin, mDia1, and profilin mediates F-actin polymerization from growing microtubule ends *in vitro* (Henty-Ridilla et al., 2016). In another example, the +TIP protein adenomatous polyposis coli (APC) was reported to promote actin assembly, activity that is negatively regulated by interaction of APC with EB1 (Juanes et al., 2020). Future work investigating how CLASP’s interactions with its binding partners may impact the crosstalk between the microtubule and actin networks is needed. It remains unclear if CLASP can interact with F-actin when targeted to growing microtubule ends via EBs, and whether such interaction could also promote actin polymerization from the dynamic microtubule ends, as shown for the complex with CLIP-170 (Henty-Ridilla et al., 2016).

Microtubule-dependent actin filament coalescence demonstrated in our minimal component system could be important for building higher-order actin structures in cells. Regulators such as APC have been shown to both mediate actin assembly and facilitate microtubule capture at focal adhesions (Juanes et al., 2017; Juanes et al., 2019; Wen et al., 2004). More recently, it has also been reported that APC can organize branched actin networks on microtubules in growth cones of hippocampal neurons (Efimova et al., 2020). In our study, we find that both actin-microtubule association and the architecture of contractile actin fibers are severely disrupted when CLASPs are depleted in vascular smooth muscle cells. Actin defects observed in actin stress fiber structures are consistent with previous findings that CLASP depletions cause defects in actin-based invasive protrusions in vascular smooth muscle cells (Efimova et al., 2014; Zhu et al., 2016). Overall, our results identify a potential role for CLASP-mediated organization of actin filaments along microtubules in the process of actin fiber assembly in contractile cells (Figure 5).

## Abbreviations used

APC: adenomatous polyposis coli
CLASP: cytoplasmic linker associated protein
F-actin: actin filaments
MAP: microtubule associated protein
PCC: Pearson correlation coefficient
+TIP: microtubule plus tip interacting protein
TIRF: total internal reflection fluorescence microscopy
TOG: tumor overexpressing gene

## ACKNOWLEDGEMENTS

We thank G. Brouhard and S. Bechstedt (McGill University) for the pHATHUGS vector and K. Grishchuk (University of Pennsylvania) for the His-EGFP-L-TOG2-S construct. We thank H. Manning and M. Lang (Vanderbilt) for MATLAB codes used for normalization of intensity line scans and extracting step information when using vbFRET software. We thank G. Arpag for the MATLAB code used for generating line scans for the stepping analysis. We thank D. Bowman and A. Olivares for help with the actin purification. Finally, we thank all members of the Zanic lab and the Vanderbilt Microtubules and Motors club for discussions and feedback. This work was supported by National Institutes of Health grant R35GM119552 to MZ. IK acknowledges support from R35GM127098. NR acknowledges support from National Institutes of Health grants F31HL151033-01A1 and T32GM0083320. EJL acknowledges the support of the National Institutes of Health IBSTO training grant T32CA119925.

## MATERIALS AND METHODS

### DNA constructs

The cDNA encoding full-length human CLASP2a was purchased from Dharmacon (Accession: BC140778.1) and subcloned into: (i) a pFastBacHT vector (Invitrogen) containing an N-terminal 6xHis-tag and; (ii) a modified pHAT vector containing an N-terminal 6xHis tag and a C-terminal eGFP and StrepII tag (a gift from S. Bechstedt and G. Brouhard, McGill University, Canada). The cDNA encoding His-EGFP-L-TOG2-S in a pRSETa vector was a gift from E. Grishchuk (University of Pennsylvania, Philadelphia, PA, USA).

### Protein biochemistry

#### His-CLASP2a

His-CLASP2α was expressed in baculovirus-infected Sf9 insect cells using the Bac-to-Bac system (Invitrogen). After the first amplification, baculovirus-infected insect cells (BIIC) stocks were used to infect Sf9 cells at a density of 1 × 106 viable cells/ml at a ratio of 10–4 BIIC:total culture volume (D. J. Wasilko et al., 2009; D. Wasilko & Lee, 2006). Cells were harvested 5 days after infection. The cell pellets were lysed by one freeze–thaw cycle and Dounce homogenizing in lysis buffer containing 50 mM PIPES (pH 6.8), 120 mM KCl, 2 mM MgCl2, 50 mM L-glutamate, 50 mM L-arginine, 10% glycerol (v/v), 0.1% (v/v) Tween-20, 1 mM DTT and supplemented with protease inhibitors. Genomic DNA was sheared by passing the lysate through an 18-gauge needle. The crude lysates were clarified by centrifugation for 20 min at 4°C and 35,000 rpm in a Beckman L90K Optima and 50.2 Ti rotor. Clarified lysates were applied to a HisTrapHP column (GE Lifesciences) according to the manufacturer’s protocol. His-CLASP2α protein was eluted in 50 mM PIPES (pH 6.8), 400 mM KCl, 5% (v/v) glycerol, 0.1% (v/v) Tween-20, 2 mM MgCl2, 1 mM DTT, 50 mM L-glutamate, 50 mM L-arginine, and a linear gradient of 50 mM - 300 mM imidazole. His-CLASP2α was then buffer exchanged into CLASP storage buffer containing 25 mM PIPES [piperazine-N,N’-bis(2-ethanesulfonic acid)] (pH 6.8), 150 mM KCl, 5% (v/v) glycerol, 0.1% (v/v) Tween-20, 50 mM L-glutamate, 50 mM L-arginine, and 1 mM DTT using an Amicon centrifugal filter.

#### His-CLASP2a-EGFP-Strep

His-CLASP2α-EGFP-Strep was expressed in Sf9 insect cells and purified as described above with the following modifications. Cell pellets were lysed in buffer containing 50 mM Tris (pH7.5), 100 mM NaCl, 5% Glycerol, 0.1% Tween-20, 1 mM DTT, supplemented with protease inhibitors. His-CLASP2α-EGFP-Strep was eluted from the HisTrap column with 50 mM HEPES pH 7.5, 150 mM NaCl, 5% glycerol, 0.1% Tween 20, 2 mM MgCl2, 1 mM DTT, 100 mM L-Glut/L-Arg, and a gradient of 50 - 500 mM imidazole. His-CLASP2α-EGFP-Strep protein was then further purified by size exclusion chromatography using a Superdex 200 Increase 10/300 GL column (Cytiva) in CLASP storage buffer.

#### His-EGFP-L-TOG2-S

His-EGFP-L-TOG2-S was expressed in BL21(DE3) E. coli cells. Expression was induced with 0.2 mM IPTG at 18°C for 16 h. Cells were lysed for 1 hr at 4°C in 50 mM HEPES (pH 7.5), 300 mM NaCl, 2 mM MgCl2, 5% (v/v) glycerol, 0.1% (v/v) Tween-20, 1 mM DTT and 40 mM imidazole and supplemented with 1 mg/ml lysozyme, 10 mg/ml PMSF and EDTA-free protease inhibitors (Roche). The crude lysate was sonicated on ice and then clarified by centrifugation for 30 min at 4°C and 35,000 rpm in a Beckman L90K Optima and 50.2 Ti rotor. Clarified lysates were applied to a HisTrapHP column (Cytiva) according to the manufacturer’s protocol. His-EGFP-L-TOG2-S protein was eluted with 50 mM HEPES (pH 7.5), 500 mM NaCl, 2 mM MgCl2, 5% (v/v) glycerol, 0.1% (v/v) Tween-20 and 1 mM DTT and linear gradient of 40 mM - 500 mM imidazole. His-EGFP-L-TOG2-S protein was then further purified by size exclusion chromatography using a Superdex 200 Increase 10/300 GL column (Cytiva) in CLASP storage buffer.

All proteins were snap frozen in single-use aliquots and protein purity was assessed by 10% SDS-PAGE (BioRad Laboratories) and stained with Coomassie Brilliant Blue (Supplemental Figure S1).

#### Tubulin purification and labeling

Tubulin was purified from bovine brains using the standard protocol (Castoldi & Popov, 2003). Briefly, tubulin was purified by cycles of polymerization and depolymerization using the high-molarity PIPES buffer method (Castoldi & Popov, 2003). Tubulin was labeled with Alexa Fluor 555 and 647 dyes (Invitrogen), and biotin (Sigma-Aldrich) following a published protocol (Hyman et al., 1991).

#### Actin purification and labeling

Actin was purified from chicken breast using the standard protocol (Pardee & Spudich, 1982). First, the chicken breast was made into a muscle acetone powder and then further purified through cycles of polymerization and depolymerization (Pardee & Spudich, 1982). Actin was labeled with Alexa Fluor 647 (Invitrogen) following a published protocol (Shekhar, 2017). Before experiments, actin was left on ice overnight and pre-clarified by spinning actin at 450,000 x g for 20 minutes at 4C in a TL Optima ultracentrifuge. Actin was stored at 4C for up to 3 weeks.

### Co-sedimentation experiments

#### High-speed co-sedimentation experiments

High-speed co-sedimentation experiments with CLASP2α were adapted from published work (Engel et al., 2014). In brief, F-actin was prepared by first pre-clarifying G-actin (Cytoskeleton, Inc.) by spinning the sample at 13,300 rpm at 4C for 15 minutes. Supernatant was collected and added to an actin polymerization buffer (50 mM KCl, 2 mM MgCl2, and 1 mM ATP) and was incubated for 1 hour at room temperature to obtain F-actin. Either 434 nM His-CLASP2α or CLASP storage buffer and 21 μM F-actin, were incubated at 37C for 30 minutes in a 100 μl reaction buffer including 100 mM NaCl, 25 mM HEPES pH 7.25, and 10 mM MgCl2. Then samples were ultracentrifuged (TL-100, Beckman Coulter) at 160,000 x g for 20 minutes at 4C. Then, 50 μL of the top supernatant was collected and 13 μL of 5X SDS Loading Dye was added. The pellet was resuspended in 63 μL of 1X SDS Loading Dye. 2 μL of 1 mg/mL bovine serum albumin (BSA, Boston BioProducts) was added to 38 μL of each top supernatant and pellet sample, boiled at 95C for 5 minutes, and then loaded onto a 10% SDS-PAGE gel for electrophoresis. Subsequently, gels were stained with Coomassie Brilliant Blue, de-stained overnight, and imaged for quantification.

#### Low-speed co-sedimentation experiments

For low-speed co-sedimentation experiments, phalloidin-stabilized F-actin was prepared by incubating 20 μM G-actin (Cytoskeleton, Inc.) with equimolar phalloidin, 50 mM KCl, and MRB80 (80 mM PIPES pH 6.8, 4 mM MgCl2, and 1 mM EDTA) on ice for 5 minutes, followed by 1 hour at room temperature. Solutions of 434 nM His-CLASP2α or 1 μM α-actinin (Cytoskeleton, Inc.) were incubated with and without 15 μM phalloidin-stabilized F-actin for 20 minutes at room temperature. Samples were subsequently centrifuged at 10,000 x g (TL-100, Beckman Coulter or accuSpin Micro 17R) for 20 minutes at 25C. Then, 50 μL of the top supernatant was collected, to avoid disturbing the pellet sample, and 13 μL of 5X SDS Loading Dye was added to the supernatant sample. The pellet was resuspended in 63 μL of 1X SDS Loading Dye. 2 μL of 1 mg/mL BSA (Boston BioProducts) was added to 38 μL of each top supernatant and pellet sample, boiled at 95C for 5 minutes, and then loaded onto a 10% SDS-PAGE gel for electrophoresis. Subsequently, gels were stained with Coomassie Brilliant Blue, de-stained overnight, and imaged for quantification.

### *In vitro* reconstitution assay conditions and imaging

#### Chamber preparation

Microscope chambers were constructed as previously described (Gell et al., 2010; Strothman et al., 2019). Channels were constructed by sandwiching 22 x 22 mm and 18 x 18 mm silanized glass coverslips together between three thin strips of parafilm. A heat block was used to melt the parafilm and stick the coverslips together. The surface was functionalized by flowing in 25 – 100 μg/ml NeutrAvidin (Thermo Scientific) for 10 minutes, followed by blocking with 1% Pluronic F127 for 30 minutes. Chambers were washed in between these steps using MRB80 or BRB80 (80 mM PIPES, 1 mM EGTA, and 1 mM MgCl2, pH 6.8).

#### Imaging

Imaging was performed using total internal reflection fluorescence (TIRF) microscopy on a Nikon Eclipse Ti microscope with a 100x/1.49 n.a. TIRF objective, equipped with Andor iXon Ultra EM-CCD camera, 488-, 561-, and 640-nm solid state lasers (Nikon Lu-NA), Finger Lakes Instruments HS-625 high-speed emission filter wheel, and standard filter sets. Laser exposure time was 100 ms for all experiments. Experiments were performed at 35C using a Tokai Hit objective heater. Images were acquired using NIS-Elements (Nikon).

#### F-actin binding along CLASP2-coated microtubule experiments

Taxol microtubules were prepared by polymerizing 28 μM tubulin (16% biotinylated and 5% Alexa647-labeled) with a microtubule polymerization mix (MRB80, 5% DMSO, 4 mM MgCl2, and 1 mM GTP). Reaction was left on ice for 5 minutes, to remove any tubulin oligomers in the stock, and then incubated for 1 hour at 37C. Then, microtubules were diluted 56 times into 10 μM Taxol (Tocris) in MRB80 (MRB80T) while in the heat block. Microtubules were then spun down in a Beckman Airfuge IM-13 at 20 psi for 5 minutes at room temperature. Pellet was resuspended in 100 μl of MRB80T. Microtubules were stored in the dark at room temperature and used within 1 week. Phalloidin-stabilized F-actin was prepared by polymerizing 3.7 μM unlabeled G-actin, stored in 2 mM Tris-HCl pH 8.0, 0.2 mM ATP, 0.5 mM DTT, 0.1 mM CaCl2, and 1 mM NaN3, with 50 mM KCl, 1 mM MgCl2, and 1 mM ATP for 1 hour at room temperature. Then, equimolar TRITC-phalloidin (Sigma-Aldrich) was added to the reaction mix and incubated in the dark for 15 minutes at room temperature. F-actin was then spun down in an airfuge at 27 psi for 10 minutes. Pellet was resuspended in MRB80, 0.2 mM ATP, and 0.5 mM DTT. F-actin was stored in 4C and used for 1 week. For control experiments with and without phalloidin stabilization, F-actin was prepared similarly as above, but was polymerized with 20% labeled A647 G-actin for an hour at room temperature, then the reaction was split into two, where one was stabilized with equimolar unlabeled phalloidin (Sigma-Aldrich), and the other half used in parallel.

Taxol microtubules were added to NeutrAvidin-functional channels to bind to the surface. Once bound after a few minutes, channel was washed with MRB80T and then washed with imaging buffer (MRB80T, 0.2 mM ATP, 40 μg/mL glucose oxidase, 40 mM glucose, 16 μg/mL catalase, 0.08 mg/mL casein, and 10 mM DTT). Then, reaction mixes with imaging buffer, and either 100 nM CLASP2α-GFP or 100 nM L-TOG2-S, and 6.5 μM TRITC-phalloidin-stabilized F-actin were added to the imaging channel while simultaneously imaging. Control experiments were performed with CLASP storage buffer. Control experiments with and without phalloidin stabilization were performed with 1 μM F-actin. All solutions were supplemented with 20 μM Taxol.

For the actin filament and microtubule co-localization experiments, CLASP2α-GFP or GFP-L-TOG2-S reaction mixes with F-actin were added to the channel and then images of all three channels (488- for CLASP protein constructs, 561- for TRITC-phalloidin F-actin, and 640- for 5% A647-labeled microtubules) were acquired every 3 seconds for 10 minutes. Several images were taken throughout the channel after the 10-minute incubation.

For F-actin accumulation experiments, a three-color image to visualize the CLASP2α-GFP (488-), TRITC-phalloidin F-actin (561-), and microtubules (640-) was taken before adding the F-actin-CLASP reaction mix as a control for fluorescence bleed through. Then, fast imaging at 5 fps for the 561-channel was started while flowing in the reaction mix to capture the initial F-actin landing events. After 5 minutes, another three-color image to visualize the CLASP (488-), F-actin (561-), and microtubule (640-) channels was taken and then 488- and 561-channels were imaged every 3 seconds for 35 minutes. Once done, a final three-color image was taken. Experiments were performed in triplicate. Control experiments with and without phalloidin were done in duplicate.

#### Actin dynamics on CLASP2-coated microtubule experiments

Taxol microtubules were prepared as described above, however were labeled with 16% biotin and 5% Alexa555 tubulin and stored in BRB80T (BRB80, 10 μM Taxol (Tocris)). After chamber preparation, Taxol microtubules were added to bind to the surface and then were washed out with BRB80T. Before adding the reaction mix, the chamber was washed with imaging buffer (BRB80, 0.1% methylcellulose, 40 μg/ml glucose oxidase, 40 mM di-glucose, 18 μg/ml catalase, 0.8 mg/ml casein, 10 mM DTT, and 0.2 mM ATP). Then, either 100 nM CLASP2α-GFP or CLASP2α-GFP storage buffer in imaging buffer was added to the chamber and incubated for 5 minutes to allow for full coating of Taxol microtubules. Then a reaction mix with 250 nM, 20% Alexa647 labeled G-actin and either 100 nM CLASP2α-GFP or CLASP2α-GFP storage buffer were added. Three-color imaging every 5 seconds for 30 minutes immediately followed. All solutions were supplemented with 20 μM Taxol. Experiments were done in triplicate.

### Cells

A7r5 rat smooth muscle cells (ATCC # CRL-1444) were grown in low-glucose (1000 mg/l) Dulbecco’s modified Eagle’s medium (DMEM) without Phenol Red, supplemented with 10% fetal bovine serum at 37°C and 5% CO2. Cells were plated on glass coverslips coated with 10 μg/ml fibronectin 24 hours prior to experiments.

#### siRNA sequences, CLASP2 expression rescue, and transfection

Two different combinations of mixed siRNA oligonucleotides against CLASP1 and CLASP2 were used. Combination 1 (custom design, Sigma) included the CLASP1-targeted siRNA sequence 5’-CGGGAUUGCAUCUUUGAAA-3’ and the CLASP2-targeted siRNA sequence 5’-CUGAUAGUGUCUGUUGGUU-3’. Combination 2 (Mimori-Kiyosue et al., 2005) included the CLASP1-targeted siRNA sequence 5’-CCUACUAAAUGUUCUGACC-3’ and the CLASP2-targeted siRNA sequence 5’-CUGUAUGUACCCAGAAUCU-3’. Non-targeting siRNA (Dharmacon) was used for controls. For siRNA oligonucleotide transfection, HiPerFect (Qiagen) was used according to the manufacturer’s protocol. Experiments were conducted 72 hours after siRNA transfection, as at this time minimal protein levels were detected.

For CLASP2 expression rescues, a GFP-labeled CLASP2 mutant using alternative codons and therefore non-silenceable by anti-CLASP2 siRNA from combination 2 (Mimori-Kiyosue et al., 2005) was transfected into depleted cells at 48 hours after siRNA transfection. For DNA transfection, Fugene6 (Roche) was used according to the manufacturer’s protocol. Cells were processed for imaging after additional 24 hours to meet the 48-hour depletion optimum.

#### Cell labeling, imaging, and image processing

For actin imaging, cells on coverslips were fixed in 4% paraformaldehyde plus 0.3% Triton X-100 in cytoskeleton buffer (10 mM MES, 150 mM NaCl, 5 mM EGTA, 5 mM glucose and 5 mM MgCl2, pH 6.1) for 10 minutes. For co-staining with tubulin, 0.1% glutaraldehyde was added into the fixative solution. The actin cytoskeleton was visualized by phalloidin conjugated to Alexa Fluor 488 (Invitrogen, Molecular Probes). Tubulin was immunostained using anti-alpha-tubulin monoclonal antibodies DM1a (Sigma-Aldrich) and Alexa 568-conjugated goat anti-mouse IgG antibodies (Invitrogen, Molecular Probes) as secondary antibodies. CLASPs were immunostained by non-paralog-specific rabbit polyclonal antibodies VU-83 (pan-CLASP antibodies) (Efimov et al., 2007). Nuclei were visualized by DAPI (ThermoFisher). Staining was performed at room temperature. After washing, samples were mounted into ProLong^®^ Gold Antifade Reagent (Invitrogen, Molecular Probes) on glass slides and stored at −20°C.

Wide-field fluorescence imaging (Figure 4 A-D, Supplemental Figure 3 A-C) was performed using a Nikon 80I microscope with a CFI APO 60× oil lens, NA 1.4 and CoolSnap ES CCD camera (Photometrics). Laser-scanning confocal imaging was performed using Nikon A1r based on a Ti-E inverted microscope with SR Apo TIRF 100× NA1.49 oil lens run by NIS Elements C software (Nikon, Tokyo, Japan). Laser scanning confocal imaging (Figure 4 E-G) was performed using a Nikon A1r with a 100x lens NA 4.5. Maximum intensity projection over the whole cell is shown in overview images. Single confocal slices processed through the emboss filter are shown in insets. In all cell images, each fluorescent channel was contrasted by whole-image histogram stretching. In overview images in Figure 4 (E-G), the tubulin channel was gamma-adjusted to highlight microtubules at the cell periphery.

#### Phenotype verification

Actin channel images (as in Figure 4A-C) were separated and coded for double-blind phenotype verification. Three researchers independently sorted images to detect actin fiber disturbance. These blinded analyses resulted in >84% of correct image designation into NT control and CLASP-depleted phenotypes.

#### CLASP Western Blotting

Western blotting was performed using the Protein Electrophoresis and Blotting System (Bio-Rad). Briefly, A7r5 cells were transfected with two different combinations of mixed siRNA oligonucleotides against CLASP1 and CLASP2 using TransIT-X2 (Mirus). After 72 h, the cells were collected, lysed and resuspended in Laemmli Sample Buffer (Bio-Rad). 20 μg total protein for each condition was resolved on 12% SDS-PAGE gels and transferred to nitrocellulose membranes (GE Healthcare Life Sciences) at 350 mA for 3 h for blotting. The membranes were then blocked with 5% nonfat dried milk (Sigma-Aldrich) for 1 h and incubated overnight with primary antibodies: anti-rat CLASP1 (KT 67, Absea), anti-rat CLASP2 (KT69, Absea) and anti-mouse GAPDH (Santa Cruz). IRDye 700 or 800 (LI-COR Biosciences) were used as secondary antibodies. The membranes were imaged on an Odyssey Infrared Imaging system (LI-COR Biosciences).

### Image Analysis

Images were processed using the Fiji (Fiji Is Just another ImageJ) image processing package (Schindelin et al., 2012).

#### Protein Gel Densitometry

Gels were quantified using the gel ImageJ plugin. Intensities were recorded and normalized to the loading control in Microsoft Excel. Normalized bands were then summed to determine percentage of protein in supernatants and pellets. Positive and negative controls were done with 2 μM α-actinin and 2 μM BSA, respectively. Plots were produced in GraphPad Prism.

#### Colocalization using Pearson Correlation Coefficient (PCC)

Three, three-color images of 488 nm- (Control, CLASP2α-GFP, or GFP-L-TOG2-S), 561 nm- (TRITC-phalloidin F-actin), and 640 nm- (5% A647-labeled microtubules) channels were cropped to 335 x 170 pixel size (~ 54 μm x 27 μm). Then, the PCC was measured using the ImageJ plugin, JACoP (Just Another Co-localization Plugin) for microtubules versus CLASP and microtubules versus F-actin (Bolte & Cordelières, 2006; Dunn et al., 2011). Individual PCC values with the standard deviation were plotted in GraphPad Prism. Kruskal-Wallis one-way ANOVA with multiple comparisons test was used to test for significance in GraphPad Prism. Experiments were done in triplicate.

#### Average Actin Intensity on CLASP2-coated Microtubules

All microscopy movies were first drift-corrected using the Image Stabilizer plugin for ImageJ. To avoid heterogeneous illumination occurring in the TIRF field, all the microscope images were cropped (~300 x 512-pixel size) to eliminate out of focus or dimmer microtubules. For F-actin landing experiments, the microtubule channel image taken between the first 5-minute movie and the second 35-minute movie was used to determine the microtubule region. For actin polymerization experiments, the last microtubule image was used. This image was thresholded, using the Auto MaxEntropy method, and recorded. Then a selection was created around the thresholded microtubule region and overlayed on the F-actin channel. A ROI was created, and the average F-actin intensity was measured using the Time Series Analyzer V3 ImageJ plugin. F-actin intensity was normalized to the maximum intensity and fit to the following intensity equation,

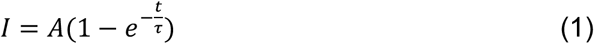

where I is the F-actin intensity, A is the maximum F-actin intensity, t is time, and τ is the half time in the MATLAB Curve Fitting Tool (MathWorks, Inc.). Results were plotted in MATLAB (MathWorks, Inc).

#### F-actin Landing Step Analysis

For kymograph production, straight or segmented lines were drawn along microtubules and then overlayed on the F-actin channel. Straight line kymographs were produced using a custom ImageJ plugin, and segmented line kymographs (used occasionally for curvy microtubules), were produced using the KymographBuilder ImageJ plugin. Kymographs were made for the first 5-minute movies and the second 35-minute movie along the same microtubule. Since F-actin landing events appear as steps, we used the open-source, vbFRET graphical user interface (gui) to measure the number of “steps” (Bronson et al., 2009). This software was created to analyze single-molecule FRET data using hidden Markov modeling and finds the most probable fit using the variational Bayesian expectation maximization algorithm (Bronson et al., 2009). For analysis, kymographs were loaded into a custom MATLAB function that extracts vertical line scans of a designated width (3-pixels) and intensity values were normalized to 90% of the maximum intensity. This normalization was recommended when analyzing non-single molecule FRET data using the vbFRET software (Bronson et al., 2009). To use this software, the FRET efficiency, which is defined as

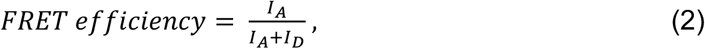

was set equal to the F-actin intensity or donor intensity (I_D_), by defining the acceptor intensity (I_A_) as,

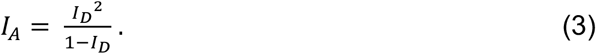

Line scans were selected based on the center-most line scan and then additional line scans were selected in intervals of 15 pixels or 2.5 microns along the microtubule lattice, as permitted by individual microtubule lengths. Selected line scans were then loaded into the vbFRET gui for stepping analysis (Bronson et al., 2009). The number of possible states was set to a minimum of 1 and maximum of 20, and the number of fitting attempts per trace was set to 25. The vbFRET session and idealized traces were saved, and steps were extracted using custom MATLAB codes. Steps were measured after F-actin was visible in solution (after 30-40 seconds). Due to overfitting of noise, step fits were filtered, with steps of sizes smaller than the 3 standard deviations below the mean step size removed. F-actin landing events followed by F-actin unbinding (negative steps) were additionally removed to consider the maximum number of F-actin accumulating on a microtubule segment. Line traces were analyzed separately for 5-minute and 35-minute movies and reported results are the summation of the two movies to ensure all F-actin landing events are measured. Example F-actin intensity trace in Figure 2, with vbFRET generated fit, was produced in MATLAB (MathWorks, Inc.) and F-actin landing event numbers were plotted in GraphPad Prism.

#### F-actin Bridge Length

After 30 minutes, F-actin bridge lengths were measured by drawing segmented lines along F-actin in ImageJ. Error bars are the standard deviation of the mean F-actin bridge lengths for each repeat. Analysis and plots were done in Microsoft Excel and GraphPad Prism.

## Notes

### Competing Interest Statement

The authors have declared no competing interest.

