## Supplemental Material for "CLASP2 facilitates dynamic actin filament organization along the microtubule lattice"

**SUPPLEMENTAL MATERIALS**

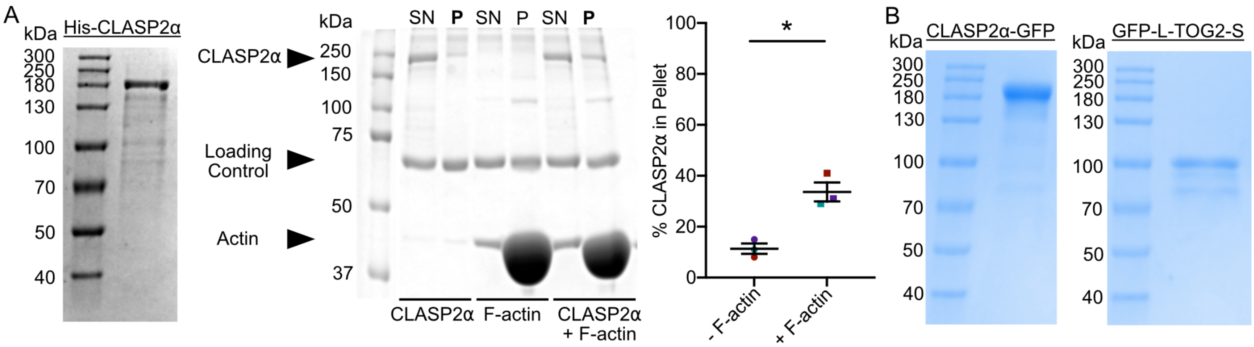


**Supplemental Figure 1.** Purified human CLASP2α directly interacts with actin filaments. A) SDS-PAGE gel showing purified His-CLASP2α. SDS-PAGE example gel for high-speed co-sedimentation with 21 μM F-actin and 434 nM CLASP2α. Quantification of high-speed co-sedimentation. Error bars represent standard error of the mean (SEM). Welch's t-test, * $p < 0.05$. Different colored data points correspond to experimental repeat. B) SDS-PAGE gels showing purified His-CLASP2α-eGFP-Strep and His-eGFP-L-TOG2-S.

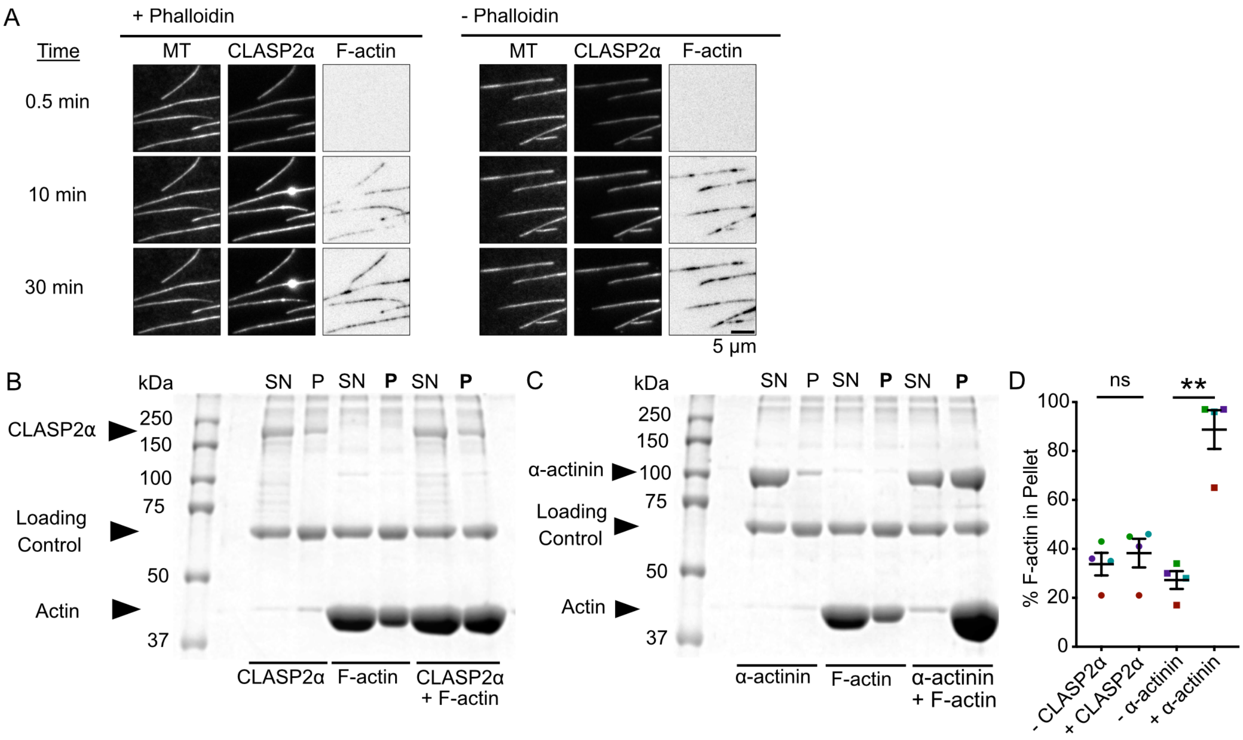


**Supplemental Figure 2.** CLASP2α does not bundle actin filaments and accumulation of actin filaments on microtubules is not dependent on TRITC-phalloidin stabilization. A) Example TIRF time-lapse images of 1 μM phalloidin-stabilized F-actin and 1 μM F-actin landing along CLASP2α-coated microtubules. Example SDS-PAGE gels from low-speed co-sedimentation experiments with CLASP2α (B) or with α-actinin (C) and F-actin. Loading control is BSA. D) Quantification of the percent of F-actin in the pellet. Error bars are the SEM. Welch's corrected t-test ns $p > 0.05$, ** $p < 0.01$. Different colored data points correspond to experimental repeat.

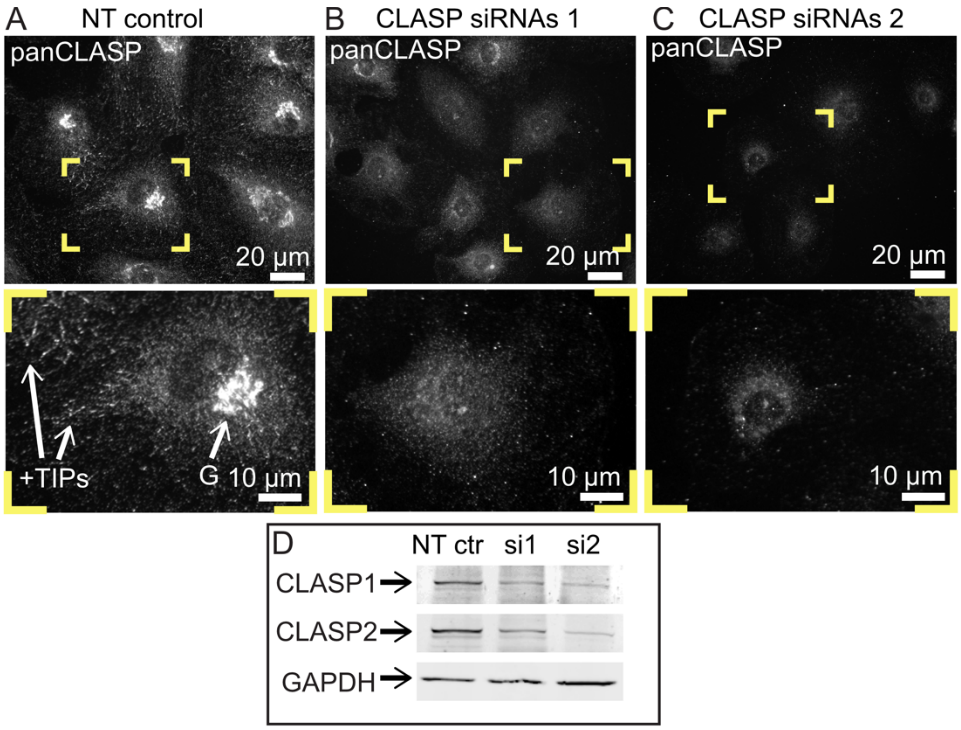


**Supplemental Figure 3.** CLASPs are efficiently knocked down by siRNA in A7r5 cells. A-C) immunostaining with pan-CLASP antibody (grayscale). A) NT control, B) siRNA combination 1, C) siRNA combination 2. Yellow boxes are enlarged below to highlight details. Scale bars, 20µm in overviews and 10µm in insets. D) Western blotting indicating CLASP1 and CLASP2 protein level reduction in cell population treated with siRNA combinations 1 and 2 as compared to control. Loading control, GAPDH.

**Video 1.** F-actin landing on CLASP2-coated microtubules. Images of Taxol microtubules (left-gray), average intensity projections of CLASP2α storage buffer (top row) or 100 nM CLASP2 (bottom row, middle-gray), time-lapse of 6.5 µM F-actin (right-black), and three-color merged time-lapse. Video play back is 100 frames per second.

**Video 2.** Dynamic actin filaments form bridges between microtubules over time. 30-minute time-lapse merged images of Taxol microtubules and actin. Video playback is 25 frames per second.
